# Parallel evolution of reduced cancer risk and tumor suppressor duplications in Xenarthra

**DOI:** 10.1101/2022.08.04.502824

**Authors:** Juan Manuel Vazquez, Maria T. Pena, Baaqeyah Muhammad, Morgan Kraft, Linda B. Adams, Vincent J. Lynch

## Abstract

The risk of developing cancer is correlated with body size and lifespan within species, but there is no correlation between cancer and either body size or lifespan between species indicating that large, long-lived species have evolved enhanced cancer protection mechanisms. Previously we showed that several large bodied Afrotherian lineages evolved reduced intrinsic cancer risk, particularly elephants and their extinct relatives (Proboscideans), coincident with pervasive duplication of tumor suppressor genes (Vazquez and Lynch 2021). Unexpectedly, we also found that Xenarthrans (sloths, armadillos, and anteaters) evolved very low intrinsic cancer risk. Here, we show that: 1) several Xenarthran lineages that independently evolved large bodies, long lifespans, and reduced intrinsic cancer risk; 2) reduced cancer risk in the stem lineages of Xenarthra and Pilosa occurred coincident with bursts of tumor suppressor gene duplications; 3) cells from sloths proliferate extremely slowly while Xenarthran cells induce apoptosis are very low levels of DNA damage; and 4) the prevalence of cancer is extremely low Xenarthrans, and cancer is nearly absent from armadillos. These data implicate the duplication of tumor suppressor genes in the evolution of remarkably large body sizes and decreased cancer risk in Xenarthrans and suggest they are a remarkably cancer resistant group of mammals.

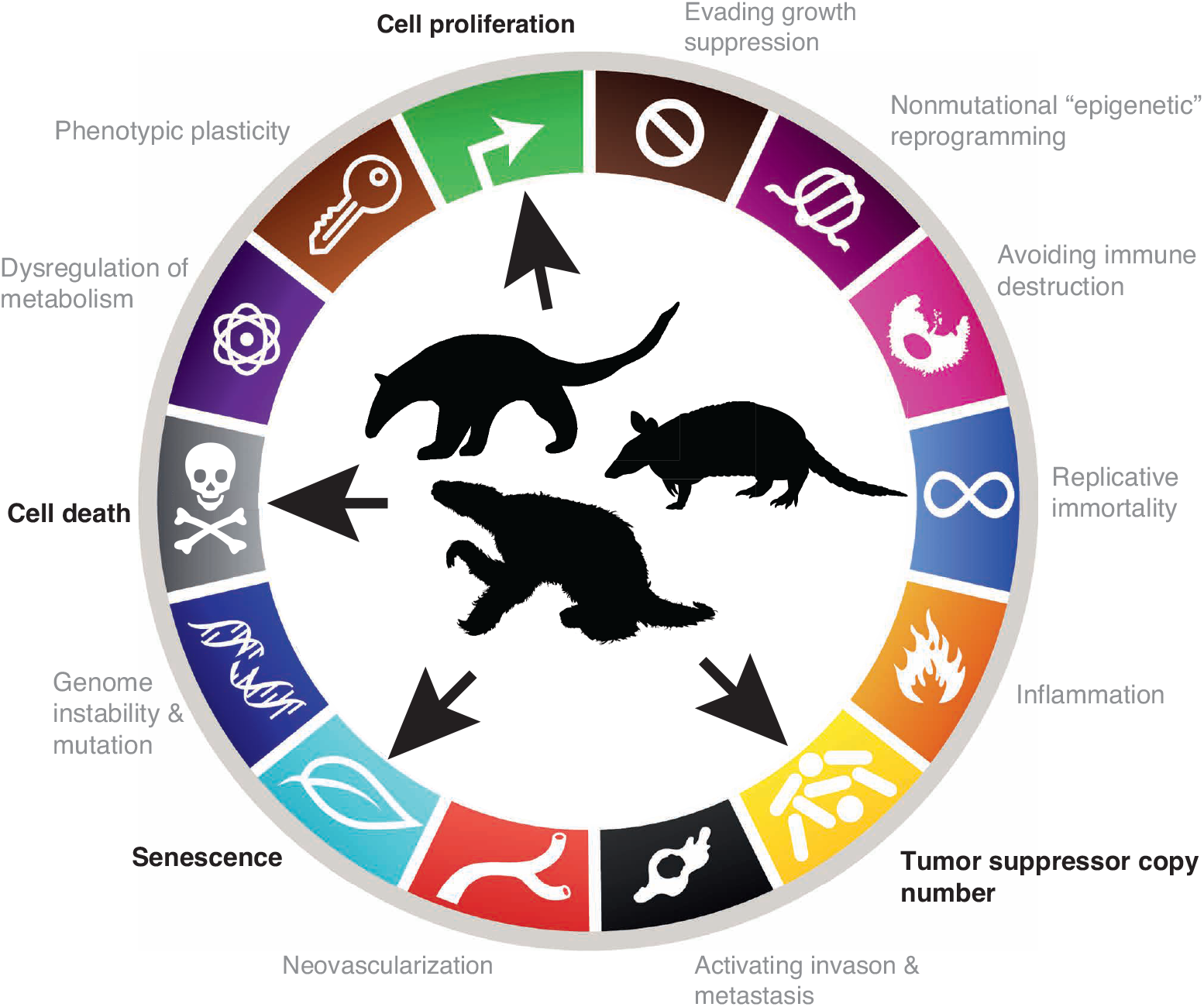

## Introduction

The evolution of very large bodies and long lifespans in animals is constrained by an increased risk of developing cancer (Caulin et al., 2015; Caulin and Maley, 2011; Nagy et al., 2007; Peto, 2015). If similar cell-types in large and small animals, for example, have a similar risk of malignant transformation and equivalent cancer suppression mechanisms, large organism with many cells should have a higher risk of developing cancer than small organisms with fewer cells; Similarly the cells of organisms with long lifespans have more time to accumulate cancer-causing mutations and other types of damage than organisms with shorter lifespans and therefore should be at an increased risk of developing cancer, a risk that is compounded in large bodied, long-lived organisms (Caulin et al., 2015; Caulin and Maley, 2011; Nagy et al., 2007; Peto, 2015). While there is a strong positive correlation between body size, age, and cancer prevalence within species (Dobson, 2013; Green et al., 2011; Nunney, 2018), there are no correlations between body size and cancer risk or lifespan and cancer risk between mammalian species (Abegglen et al., 2015; Boddy et al., 2020; Bulls et al., 2022; Vincze et al., 2022) – this lack of correlation is often referred to as ‘Peto’s Paradox’ (Peto, 2015).

Many mechanisms can potentially resolve Peto’s paradox, but they have only been experimentally studied in a few species, such as rodents (Azpurua and Seluanov, 2013; Gorbunova et al., 2012; Liang et al., 2010; Salmon and Akha, 2008; Seluanov et al., 2009; Tian et al., 2019; Zhang et al., 2021), bats (Foley et al., 2018; Kacprzyk et al., 2021; Koh et al., 2019), turtles (Glaberman et al., 2021), elephants (Abegglen et al., 2015; Sulak et al., 2016; Vazquez et al., 2018), and even Drosophila (Garschall et al., 2017; Parkes et al., 1998; Peleg et al., 2016; Shepherd et al., 1989). We and others have shown that elephants, for example, evolved cells that are extremely sensitive to DNA damage (Abegglen et al., 2015; Sulak et al., 2016) at least in part through duplication of tumor suppressor genes (Caulin et al., 2015; Sulak et al., 2016; Tollis et al., 2020; Vazquez et al., 2018; Vazquez and Lynch, 2021); While this burst of tumor suppressor duplication occurred coincident with the evolution of reduced intrinsic cancer risk in Proboscideans (Vazquez and Lynch, 2021), we found that some other mammalian lineages such as the Xenarthra (armadillos, sloths, and anteaters) (**Figure 1A**) may also have evolved reduced intrinsic cancer risk and increased tumor suppressor dosage (Vazquez and Lynch, 2021). Interestingly while living Xenarthrans are relatively small bodied, several lineages of recently extinct sloths (*Megatherium* and *Mylodon*) and armadillos (*Glyptodon*) independently evolved very large body sizes (**Figure 1B**) (Delsuc et al., 2016) indicating that at least some Xenarthrans have the developmental potential to be much larger bodied than extant species and thus must have evolved ways to reduce their intrinsic cancer risk.

**Figure 1.**
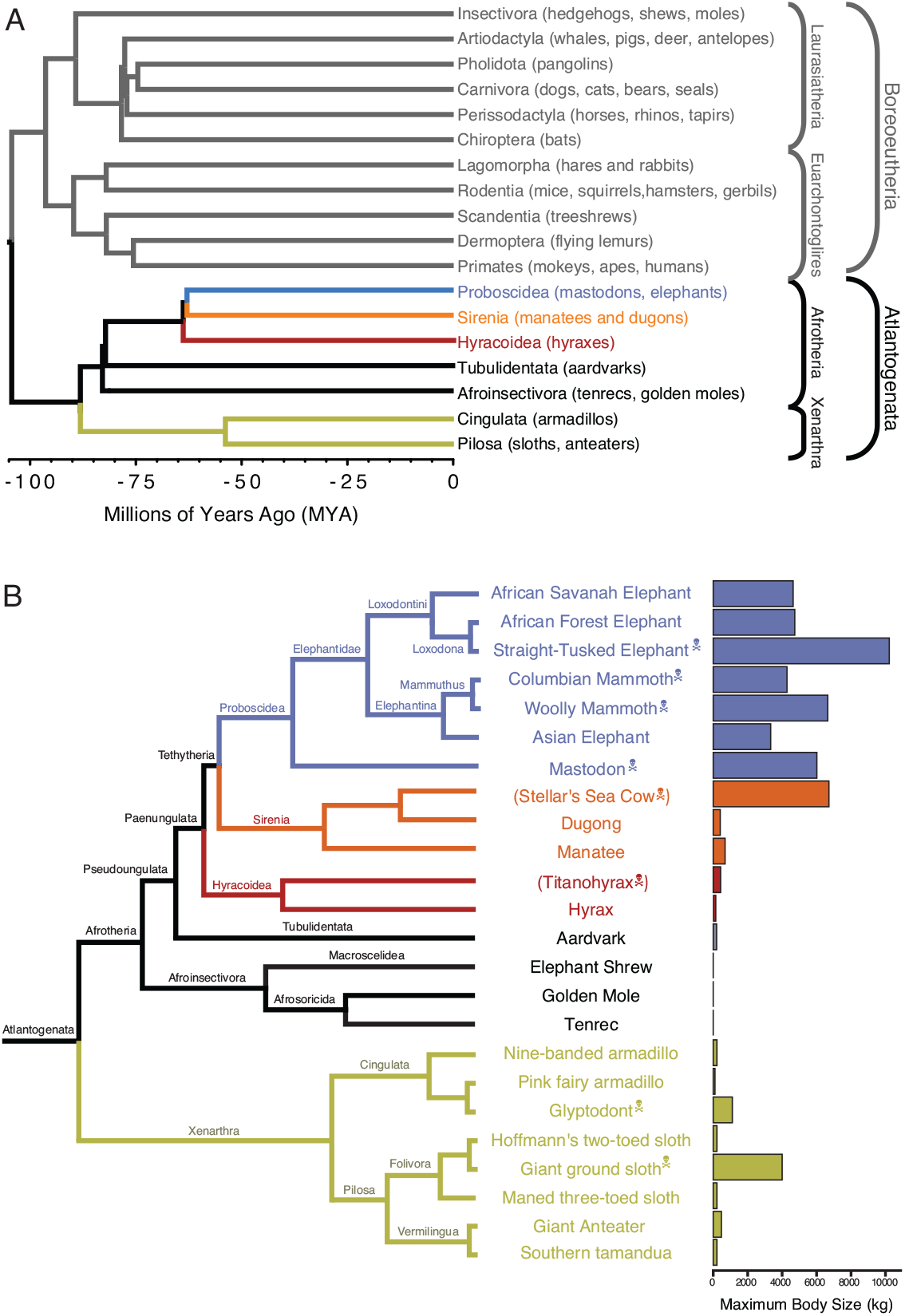
Eutherian phylogenetic relationships. **(A)** Phylogenetic relationships between Eutherian orders, examples of each order are given in parenthesis. Horizontal branch lengths are proportional to time since divergence between lineages (see scale, Millions of Ago (MYA)). The clades Atlantogenata and Boreoeutheria are indicated, the order Proboscidea is colored blue, Sirenia is colored orange, and Hyracoidea is colored red. **(B)** Phylogenetic relationships of extant and recently extinct Atlantogenatans with available genomes are shown along with clade names and maximum body sizes. Note that horizontal branch lengths are arbitrary, species indicated with skull and crossbones are extinct and do not have available nuclear genomes.

Here we reconstruct the evolution of body mass and intrinsic cancer risk across living and extinct Xenarthrans and show that intrinsic cancer risk decreased dramatically in the Xenarthran stem-lineage as well as several extinct lineages of giant armadillos and sloths. Using comparative genomics and phylogenetic methods we found that these episodes of decreased cancer risk occurred coincident with the bursts of tumor suppressor gene duplication in the Xenarthran and Pilosan stem-lineages and the evolution of cells that are particularly sensitive to DNA damage. Finally, we compare cancer prevalence across species using a new dataset of pathology reports from a large colony nine-banded armadillos (*Dasypus novemcinctus*) and show that the prevalence of cancer is lower in Xenarthrans than most other mammalian lineages and that armadillos have among the lowest cancer prevalence reported for any mammal. Similar to our previous study of body size evolution in Proboscideas, these data suggest that duplication of tumor suppressor genes occurred coincident with reductions in intrinsic cancer risk and likely facilitated the evolution of several extinct species with exceptionally large body sizes and low cancer risk.

## Results and Discussion

### Evolution reduced cancer risk in Xenarthra

We previously used a dataset of lifespan and body size data from over 1600 Eutherian mammals, focusing on Afrotheria, and used ancestral reconstruction to identify lineages to show that substantial accelerations in the rate of body mass and lifespan evolution occurred in multiple lineages, most notably in large bodied Afrotherians (**Figure 2**) (Vazquez and Lynch, 2021). Unexpectedly, we also observed that several Xenarthran lineages also showed episodes of rapid body size and lifespan evolution suggesting this clade may also be a good representative in which to study the relationships between life history evolution and cancer risk (Vazquez and Lynch, 2021). To explore the relationships between body size, lifespan, and intrinsic cancer risk in Xenarthrans in greater detail we assembled a time-calibrated supertree of Eutherian mammals by combining the time-calibrated molecular phylogeny of Bininda-Emonds *et al*. (Bininda-Emonds et al., 2007), the time-calibrated total evidence Afrotherian phylogeny from Puttick and Thomas (Puttick and Thomas, 2015) and the time-calibrated Xenarthran phylogeny of Delsuc *et al*. (Delsuc et al., 2016). While the Bininda-Emonds *et al*. phylogeny includes 1,679 species, no fossil data are included. The inclusion of fossil data from extinct species is essential to ensure that ancestral state reconstructions of body mass are not biased by only including extant species. This can lead to inaccurate reconstructions, for example, if lineages convergently evolved larger body masses from a small-bodied ancestor. In contrast the total evidence Afrotherian phylogeny of Puttick and Thomas includes 77 extant species and fossil data from 39 extinct species, while the Delsuc *et al*. Xenarthran phylogeny is based on mitogenomes for 25 living and 12 extinct Xenarthran species that are broadly representative of the Xenarthran diversity; The combined supertree includes body size data for 1775 Eutherian mammals.

**Figure 2.**
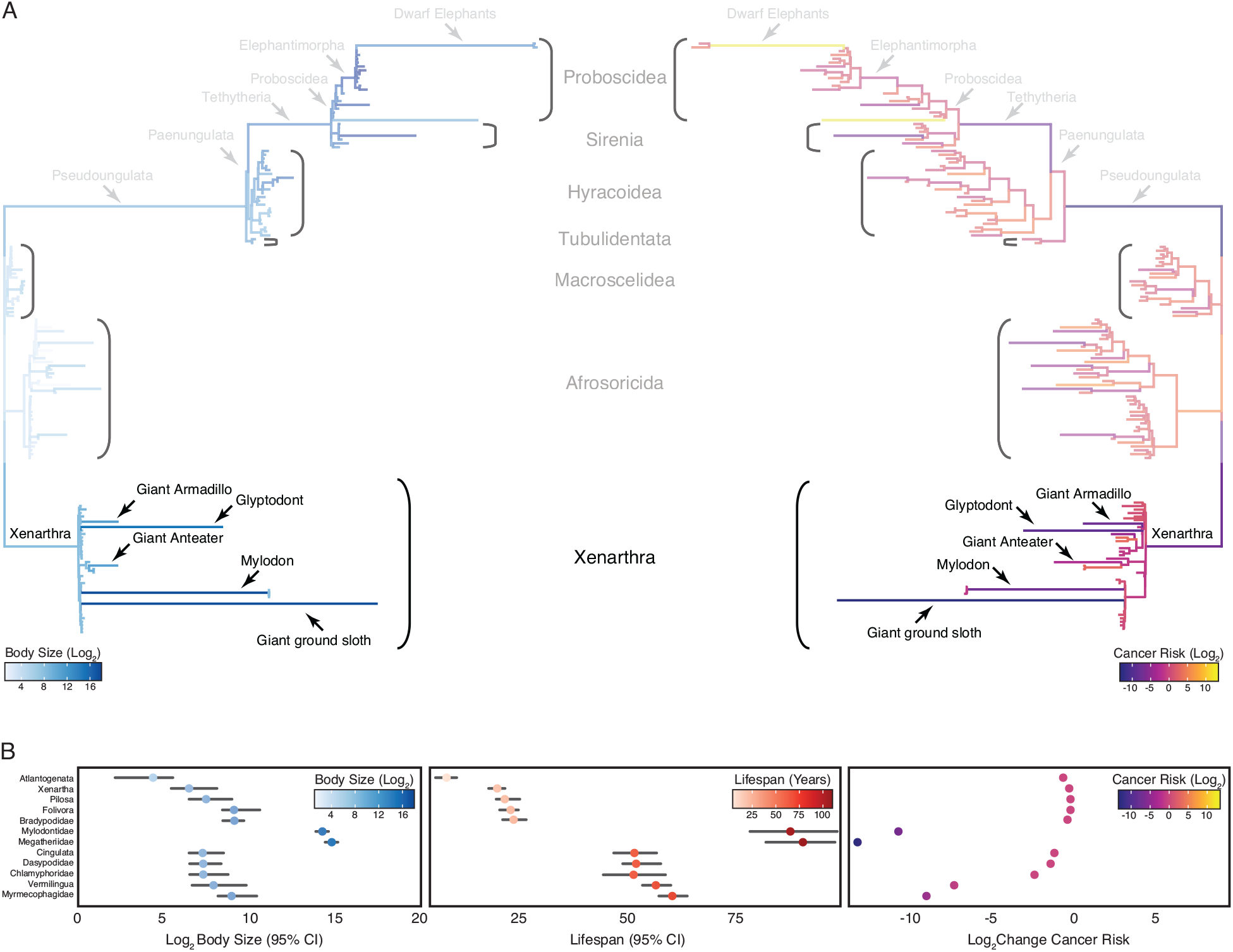
Convergent evolution of large-bodied, cancer resistant Xenarthrans. **(A)** Atlantogenata phylogeny, with branch lengths scaled by log_2_ change in body size (left) or log_2_ change in intrinsic cancer risk (right). Branches are colored according to ancestral state reconstruction of body mass or estimated intrinsic cancer risk. The Afrotherian part of the tree is shown opaque because it was analyzed in Vazquez and Lynch (2021). **(B)** Ancestral reconstructions of body size (left), lifespan (middle), and change in intrinsic cancer risk. Data are shown as mean (dot) along with 95% confidence interval (CI, whiskers) for body size and lifespan (which is estimated from body size data).

Next, we jointly estimated rates of body mass evolution and reconstructed ancestral states using the “Stable Model” implemented in StableTraits (Elliot and Mooers, 2014), which allows for large jumps in traits and has previously been shown to out-perform other models of body mass evolution, including standard Brownian motion models, Ornstein–Uhlenbeck models, early burst maximum likelihood models, and heterogeneous multi-rate models (Elliot and Mooers, 2014). Finally, we used extant and reconstructed body mass and lifespan data to estimate intrinsic cancer risk (*K*) via a simplified multistage cancer risk model: *K* ≈ *Dt*^6^ where *D* is maximum body size and *t* is the maximum lifespan across Xenarthran lineages. As expected, we found that body mass and lifespan data were correlated, that large-bodied lineages tended to also be long-lived, and that intrinsic cancer risk in Xenarthra also varies with changes in body size and longevity, with particularly notable decreases in intrinsic cancer risk in the Xenarthran stem-lineage, giant armadillo, Glyptodont, giant anteater, Mylodon, and Giant ground sloth (**Figure 2**).

### Pervasive duplication of tumor suppressors in Xenarthra and Pilosa

We previously found that reductions in intrinsic cancer risk in Afrotherians, particularly in the stem-lineage of Proboscideans, was associated with pervasive duplication of genes in tumor suppressor pathways (Vazquez and Lynch, 2021). To test if duplication of tumor suppressors was similarly common in Xenarthran lineages with reduced cancer risk, we identified duplicated genes in the genomes Afrotherians (*Orycteropus afer, Trichechus manatus, Procavia capensis*, and *Loxodonta africana*) and Xenarthrans, including three sloths (*Bradypus variegatus, Choloepus didactylus, Choloepus hoffmanni*), three armadillos (*Chaetophractus vellerosus, Dasypus novemcinctus*, and *Tolypeutes matacus*), and two anteaters (*Myrmecophaga tridactyla* and *Tamandua tetradactyla*), inferred the lineage(s) in which each duplication occurred using maximum likelihood based ancestral state reconstruction implemented in IQ-TREE2 (Minh et al., 2020), and tested if duplicate genes were enriched in Reactome pathways (Jassal et al., 2020) using the hypergeometric test implemented in WebGestalt (Liao et al., 2019). We thus identified 149 duplicates in the Xenarthran stem-lineage, 139 duplicates in the Cingulate stem-lineage, and 147 duplicates in the Pilosan stem-lineage; very few duplicate genes were identified in other Xenarthran lineages (**Figure 3A**). Duplicates in the Xenarthran stem-lineage were enriched in 92 pathways (FDR q-value ≤ 0.25), of which 26 (28%) were related to cancer biology such as cell cycle regulation, protein folding, intrinsic apoptosis, and regulation of p53 degradation while duplicates in the Pilosan stem-lineage were enriched in 13 pathways (FDR q-value ≤ 0.25), of which 8 (61%) were related to cancer biology all of which involved the cell cycle (**Figure 3B**). In contrast, duplicates in the Cingulate stem-lineage were only enriched in a single pathway (FDR q-value ≤ 0.25), “Cholesterol biosynthesis” (enrichment ratio = 15.87, hypergeometric p=1.00×10^−4,^ FDR q=0.18) which appears to be unrelated to cancer biology in a direct way unlike Xenarthran and Pilosan duplicates. These observations are consistent with earlier studies suggesting that sloths and armadillos have extra copies of tumor suppressor genes (Tollis et al., 2020; Vazquez and Lynch, 2021)

**Figure 3.**
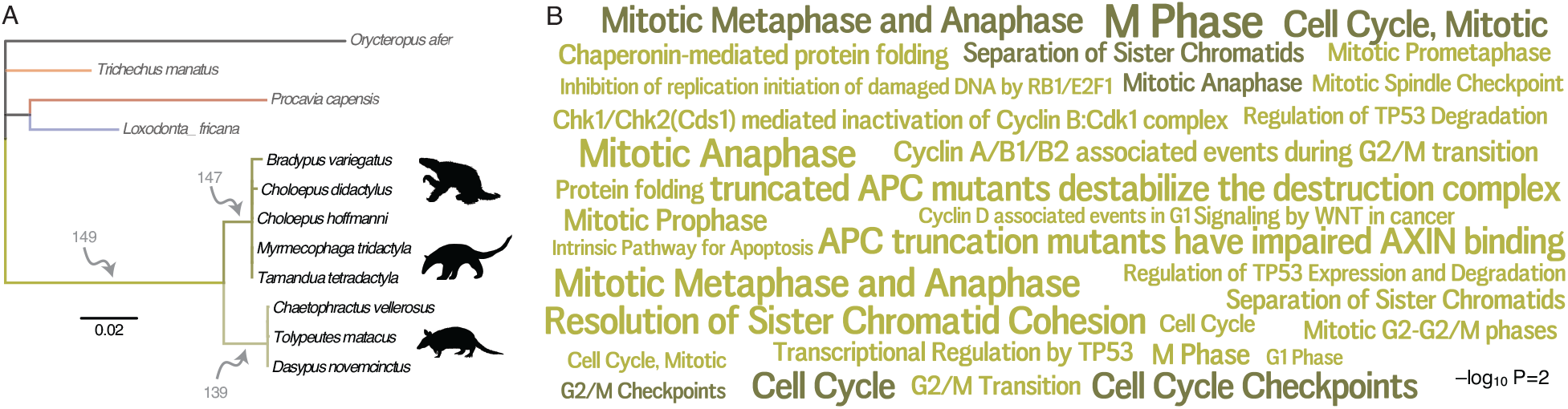
Pervasive duplication of tumor suppressors in Xenarthra. **(A)** Atlantogenata phylogeny indicating the number of genes duplicated in each Xenarthran lineage inferred by maximum likelihood. Branch lengths are drawn proportional to the per gene duplication rate (see inset scale) and colored according to lineage relationships shown in figure 1. **(B)** Word cloud of Reactome pathways in which Xenarthan (green) and Pilsoan (dark green) gene duplicates are enriched (FDR q≤0.25). Pathway term sizes are scaled according to the –log_10_ hypergeometric p-value of their enrichment (see inset scale). **Figure 3 – source data 1**. Genes duplicated in the stem-lineage of Xenarthra. **Figure 3 – source data 2**. Genes duplicated in the stem-lineage of Pilosa. **Figure 3 – source data 3**. Genes duplicated in the stem-lineage of Cingulata. **Figure 3 – source data 4**. Pathways in which genes duplicated in the stem-lineage of Xenarthra are enriched. **Figure 3 – source data 5**. Pathways in which genes duplicated in the stem-lineage of Pilosa are enriched **Figure 3 – source data 6**. Pathways in which genes duplicated in the stem-lineage of Cingulata are enriched.

### Sloth cells have extremely long doubling times

The enrichment of gene Xenarthran and Pisolan duplications in pathways related to the cell cycle suggest that cells from these species may have different cell cycle dynamics than other species. Unfortunately, measurements of cell cycle phase length or overall duration are not available for most species, however, many studies report population doubling times from mammalian cells in culture; population doubling time is proportional to the cumulative length of individual cell cycle phases, particularly the length of G_1_ (Blank et al., 2018). Therefore we performed a thorough literature review and assembled a dataset of population doubling times from 76 species representing all mammalian orders (**Figure 4A – source data 1**) and directly estimated population doubling time for primary fibroblasts from Southern three-banded armadillo (*Tolypeutes matacus*), screaming hairy armadillo (*Chaetophractus vellerosus*), six-banded armadillo (*Euphractus sexcinctus*), Northern tamandua (*Tamandua mexicana*), Southern tamandua (*Tamandua tetradactyla tetradactyla*), Linnaeus’s two-toed sloth (*Choloepus didactylus*), and Hoffmann’s two-toed sloth (*Choloepus hoffmanni*).

**Figure 4.**
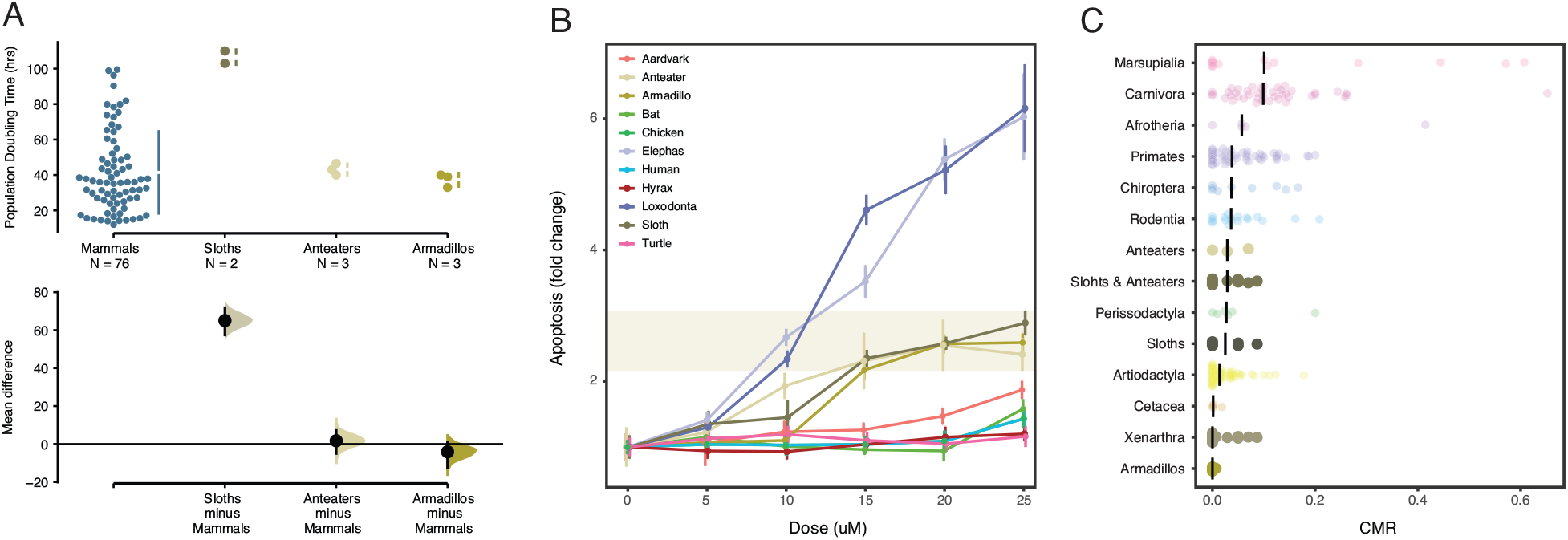
Xenarthrans have anti-cancer cellular phenotypes and a low prevalence of cancer. **(A)** Population doubling time estimates for mammalian cells (upper) and mean differences (lower). Mean differences are depicted as a dot with 95% confidence intervals as vertical error bars; P values are reported from a two-sided permutation t-test with 5000 reshuffles of control and test labels. The unpaired mean difference between Mammals and Sloths is 65.1 (57.5 - 71.8), P=4.0×10^_-4_^. The unpaired mean difference between Mammals and Anteaters is 1.71 (5.0 - 7.16), P=0.905. The unpaired mean difference between Mammals and Armadillos is −4.1 (12.5 - 0.86), P=0.781. **(B)** Induction in apoptosis in vertebrate cells by mitomycin C. Data are shown as fold change in caspase 3/7 activity (apoptosis), mean (dot) and standard deviation (vertical line); n=12 for each datapoint. **(C)** Cancer prevalence data for each species (dot) grouped by clade and shown as jittered stripchart. The number of species per clade is: Afrotheria=5, Artiodactyla=68, Carnivora=43, Cetacea=4, Chiroptera=9, Marsupialia=13, Perissodactyla=6, Primates=47, Rodentia=18, anteaters=3, sloths=6, armadillos=5, and Xenarthra=14. Medians for each clade are shown with vertical black bars. **Figure 4 – source data 1**. Data plotted in Figure 4A. **Figure 4 – source data 2**. Data plotted in Figure 4B. **Figure 4 – source data 3**. Data plotted in Figure 4C.

While there was considerable variation in population doubling times between mammalian cells, armadillo (unpaired mean difference = −4.1, two-sided permutation t-test P-value=0.09) and anteater (unpaired mean difference = 1.7, two-sided permutation t-test P-value=0.78) cells had similar doubling times as other mammals (**Figure 4A**). Sloth cells, however, had significantly longer doubling times than cell lines from other all other species with an unpaired mean difference in doubling time of 65.1 hours (95.0% CI 57.5-71.8) and a two-sided permutation t-test P-value=4.0×10^−4^ (**Figure 4A**). Our doubling time estimates for Hoffmann’s two-toed sloth (103 hrs) is similar to a previously published estimate for this species (107 hrs), and our doubling time estimates Northern tamandua (43 hrs) and Southern tamandua (40 hrs) are similar to a previously published estimate (46 hrs) for the giant anteater (*Myrmecophaga tridactyla*) suggesting these doubling times are reliable (Gomes et al., 2011). Thus we conclude that sloth cells have the longest reported doubling time for any mammal, which may allow for more time to correct DNA damage and other genomic instabilities such as aneuploidy. However, we note that doubling times may reflect biological differences in cell cycle length between species as well as technical differences that are not directly related to species-specific differences in doubling time or cell cycle length.

### Xenarthran cells are very sensitive to DNA damage

Our observation that duplicate genes in the Xenarthran stem-lineage are enriched in tumor suppressor pathways, including many pathways related to cell cycle regulation as well as “Intrinsic Pathway for Apoptosis” (enrichment ratio = 6.76, hypergeometric P=9.75×10^−3^, FDR q=0.20), suggests that Xenarthran cells may respond to genotoxic stresses differently than other species. Indeed, we previously compared the sensitivity of African elephant, Asian elephant, South African rock hyrax, East African aardvark, and Southern Three-banded armadillo primary dermal fibroblasts to DNA damage induced with mitomycin C, doxorubicin, and UV-C and found that while elephant cells had the greatest apoptotic response, Southern Three-banded armadillo cells had a greater apoptotic response than aardvark and hyrax cells but less than elephant cells (Sulak et al., 2016); This previous study was focused on the DNA damage response in elephant cells and only included a single Xenarthran, thus we could not determine if the behavior of Southern Three-banded armadillo cells was typical for Xenarthrans. To explore this observation in greater detail we treated African elephant, Asian elephant, South African rock hyrax, East African aardvark, and Southern Three-banded armadillo, Southern tamandua, Hoffmann’s two-toed sloth, human, little brown bat, chicken, and turtle primary dermal fibroblasts with increasing doses of mitomycin C (MMC), which induces multiple types of DNA damage (Lee et al., 2006), and assayed the induction of apoptosis using an ApoTox-Glo assay. Consistent with our previous study (Sulak et al., 2016), we found that Southern Three-banded armadillo, Southern tamandua and Hoffmann’s two-toed sloth cells induced apoptosis at lower MMC doses and had a greater response effect size than South African rock hyrax, East African aardvark, human, bat, chicken and turtle cells, but less than African and Asian elephant cells. These data indicate that Xenarthran fibroblasts are particularly sensitive to DNA damage.

### Extremely low cancer prevalence in Xenarthrans, especially armadillos

Previous studies have reported sporadic cases of cancer in sloths (Amaral et al., 2022; Linnehan et al., 2019; Salas et al., 2014) and anteaters (Madsen et al., 2017; Sanches et al., 2013) but there have been few systematic studies of cancer prevalence in Xenarthra. Remarkably, however, Vincze et al. (2022) found only one case of neoplasia in 22 necropsies of Linnaeus’s two-toed sloth (*Choloepus didactylus*) suggesting that cancer is uncommon in at least one sloth species. To explore cancer prevalence in Xenarthra in greater detail, we gathered published mortality data from Pilosa, including giant anteater (*Myrmecophaga tridactyla*), southern tamandua (*Tamandua tetradactyla*), silky anteater (*Cyclopes* sp.), maned three-toed sloths (*Bradypus torquatus*), brown-throated sloths (*Bradypus variegatus*), pale-throated sloth (*Bradypus tridactylus*), Linné’s two-toed sloth (*Choloepus didactylus*), as well three-toed sloths (*Bradypus sp*.) and two-toed slots (*Choloepus sp*.) without species level identification, from four published surveys (Arenales et al., 2020a, 2020b; Diniz and Oliveira, 1999; Diniz et al., 1995). These data indicate that neoplasia prevalence was only 1.8% (95% CI: 0.02 – 6.5%) in sloths, 2.3% (95% CI: 0.08 – 5.7%) in anteaters, and 2.2% (95% CI: 0.09 – 4.6%) in Pilosa (**Table 1**).

**Table 1.** Cancer prevalence in Xenarthrans. Clopper-Pearson exact 95% CIs are reported for each taxa, asterisks (*) indicate taxa with more than 82 pathology reports. Data from more than two studies are indicated with “(combined)”. **Table 1 – source data 1. Vital statistics for armadillos in the NHDP colony 2010-2020**

| Common name | Species name | Necropsies | Neoplasia | Prevalence (95% CI) |
| --- | --- | --- | --- | --- |
| Southern tamandua (combined) | *Tamandua tetradactyla* | 60 | 1 | 0.07 (0.0004–0.089) |
| Giant anteater (combined)* | *Myrmecophaga tridactyla* | 139 | 4 | 0.029 (0.008–0.072) |
| Silky anteater | *Cyclopes sp.* | 3 | 0 | 0 (0–0.708) |
| Maned three-toed sloth | *Bradypus torquatus* | 25 | 0 | 0 (0–0.137) |
| Pale-throated sloth | *Bradypus tridactylus* | 2 | 0 | 0 (0–0.841) |
| Brown-throated sloth | *Bradypus variegatus* | 8 | 0 | 0 (0–0.369) |
| Three-toed sloths (combined) | *Bradypus sp.* | 69 | 0 | 0 (0–0.050) |
| Linné’s two-toed sloth (combined) | *Choloepus didactylus* | 23 | 2 | 0.087 (0.011–0.28) |
| Two-toed sloths (combined) | *Choloepus sp.* | 40 | 2 | 0.050 (0.006–0.169) |
| Nine-banded armadillo (combined)* | *Dasypus novemcinctus* | 275 | 2 | 0.007 (0.0009–0.026) |
| Six-banded armadillo | *Euphractus sexcinctus* | 48 | 0 | 0 (0–0.074) |
| Naked-tailed armadillos | *Cabassous sp.* | 5 | 0 | 0 (0–0.522) |
| Three-banded armadillo | *Tolypeutes sp.* | 4 | 0 | 0 (0–0.602) |
| Giant armadillo | *Priodontes maximus* | 1 | 0 | 0 (0–0.975) |
| Vermilingua (anteaters)* |  | 202 | 5 | 0.023 (0.008–0.057) |
| Folivora (sloths)* |  | 109 | 2 | 0.018 (0.002–0.065) |
| Pilosa (sloths and anteaters)* |  | 311 | 7 | 0.022 (0.009–0.046) |
| Cingulata (armadillos)* |  | 333 | 2 | 0.006 (0.0007–0.021) |
| Xenarthra (sloths, anteaters, armadillos)* |  | 644 | 9 | 0.014 (0.006–0.026) |

Similarly cancer has been reported in nine-banded armadillos (Lee et al., 2015; Madsen et al., 2017; Pence et al., 1983), including a thalidomide induced malignant choriocarcinoma that perforated the uterus and metastasized to the liver, spleen, mesentery, and lungs (Marin-Padilla and Benirschke, 1963). But in the only study to systematically evaluate cancer prevalence, Boddy et al. (2020) found only two cases of neoplasia (leiomyoma and bronchial adenoma) in 69 necropsies of nine-banded armadillos (*Dasypus novemcinctus*) suggesting cancer is rare in armadillos. Therefore we explored neoplasia prevalence in armadillos from published mortality data and found that no neoplasias were reported from 55 nine-banded armadillos, 48 six-banded armadillos (*Euphractus sexcinctus*), five greater naked-tailed armadillos (*Cabassous sp*.), four three-banded armadillos without species level identification (*Tolypeutes sp*.), or one giant armadillo (*Priodontes maximus*) (Diniz et al., 1997). This remarkably low neoplasia prevalence in armadillos prompted us to compile neoplasia prevalence in a large colony of wild caught nine-banded armadillos with extensive post-mortem veterinary pathology reports managed by the National Hansen’s Disease Program (NHDP) Laboratories. We surveyed 153 pathology reports from *Dasypus novemcinctus* from 2010 to 2020 and found no cases of neoplasia. When combined with neoplasia prevalence data from Boddy et al. (2020) and Diniz et al. (1997), the estimated cancer prevalence in *Dasypus novemcinctus* was only 0.73% (95% CI: 0.09 – 2.6%) in 275 individuals (**Table 1**).

Next, we compared neoplasia prevalence in Xenarthra to other Therian mammals from two published studies that included pathology reports from 37 (Boddy et al., 2020) and 191 (Vincze et al., 2022) species of Therian mammals, as well as three species of cetaceans (Albuquerque et al., 2018); The total dataset includes cancer prevalence data from 221 species. As a group the mean prevalence of neoplasia in Xenarthra was 1.4% (95% CI: 0.6 – 2.7%), in contrast the mean prevalence of neoplasia in across other Eutherian mammals was 17.57% (95% CI: 14.86 – 20.53%) but there was much variation across species and orders (**Figure 4**). Cancer prevalence was lowest among armadillos (Cingulata) and Xenarthra (**Figure 4**). Thus, we conclude that Xenarthrans, and especially armadillos, are particularly cancer resistant mammals.

### Caveats and limitations

This study has several inherent limitations. First, genome quality can play an important role in our ability to identify duplicate genes and most Xenarthran species lack high quality reference genomes (**Figure 5**) or transcriptome annotations making assessment of functional gene duplication difficult for some species (Vazquez and Lynch, 2021). We also assume that gene duplicates maintain ancestral tumor suppressor functions and increase cancer resistance through dosage effects or by providing redundancy to loss of function mutations but sub- and neofunctionalization between paralogs can lead to functional divergence between paralogs (Innan and Kondrashov, 2010) and developmental systems drift (Lynch, 2009; True and Haag, 2001) can lead to functional divergence between orthologs and incorrect pathway assignment (Altenhoff et al., 2012; Liao and Zhang, 2008; Stamboulian et al., 2020). Second, while we have shown that fibroblasts from Xenarthra are particularly sensitive to DNA damage, and that sloths cells have very long doubling times, these studies used only a single cell-type from a single individual but there is likely variation in these traits across cell-types and individuals; We also have not shown that duplicate genes contribute to these cellular phenotype. However, our observation that cells from different vertebrate classes and mammalian orders have similar apoptotic responses to MMC and doubling times suggests that species effects are greater than possible effects from individual variation.

**Figure 5.**
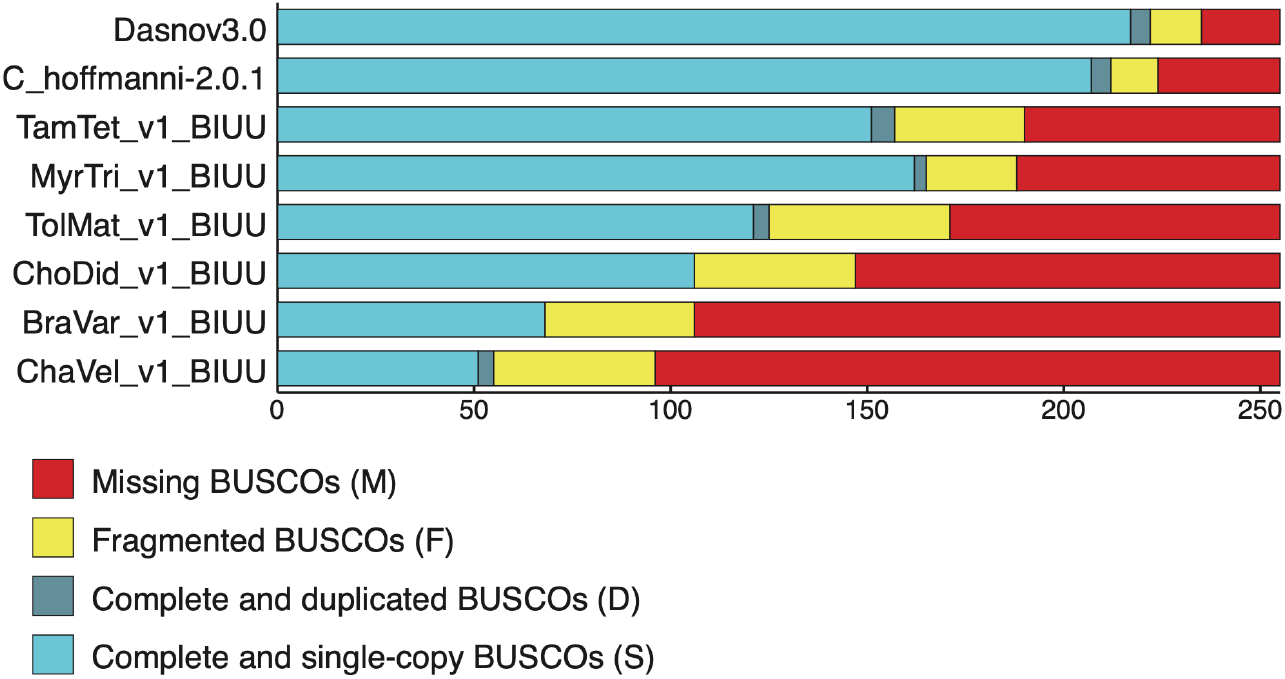
BUSCO scores for Xenarthran genomes used to estimate gene copy number.

The estimation of cancer prevalence can be error-prone particularly for species without long term morbidity and mortality studies. The probability of detecting at least one individual with cancer, for example, is strongly dependent on sample size; Indeed, cancer was detected in at least one individual in almost all species with more than 82 individual pathological records illustrating that cancer is likely to be detected in all mammals with adequate sample sizes (Vincze et al., 2022). The number of post-mortem pathology reports we gathered from the literature for Xenarthrans was below 82 for all species except giant anteater and nine-banded armadillo suggesting caution when interpreting the low cancer prevalence for other species. However our dataset included 275 pathology reports from *Dasypus novemcinctus*, of which only two had reports of neoplasia and no reports of natural metastasis, indicating that the low prevalence of cancer in *Dasypus novemcinctus* is reliable. Finally while cancer prevalence from 333 armadillos is extremely low at 0.6% (95% CI: 0.07 – 2.1%), our dataset only includes armadillo species from the family Dasypodidae and additional data from the other armadillo family (Chlamyphoridae) is necessary to determine if this low cancer prevalence is a common trait of armadillos (Cingulata) or limited to Dasypodidae.

### Ideas and speculation

Extant Xenarthrans are relatively small bodied, for example, sloths generally weigh 3.5 - 5.5 kg, armadillos 120 g - 33 kg, and tamanduas and anteaters 400 g-28.5 kg. In stark contrast, recently extinct sloth and armadillo species that lived during the pleistocene were much larger – the extinct armadillo *Glyptodon* is estimated to have been roughly the size of a Volkswagen Beetle and weighed 800–840 kg whereas the ground sloth *Megatherium americanum* is one of the largest known land mammals weighing nearly 4,000 kg and measuring up to 6 m in length (Pant et al., 2014). Remarkably, a previous study found that the rate of body size evolution in Megatheriidae was very rapidly, perhaps as fast as 129 kg/My (95% CI: 42 – 384 kg/My), while the ∼5.5 kg *Choloepus* was deeply nested within a clade with an average body mass of 236 kg (Pant et al., 2014). These data indicate that the living Xenarthrans represent a biased sample of Xenarthran body sizes throughout history, particularly sloths and armadillos, and have the developmental potential to rapidly evolve large bodies. Thus, they must also have evolved mechanisms that reduce their cancer risk. Our data suggests that a burst of tumor suppressor duplication in the Xenarthran and Pilosan stem-lineages may have contributed to the evolution of this reduced cancer risk at least in part by lowering the sensitivity of Xenarthran cells to DNA damage which prevent the accumulation of mutations that give rise to cancer as a byproduct of DNA repair. While in Vazquez and Lynch (2021), the availability of ancient elephantid genomes enabled the resolution of body size and tumor suppression copy number co-evolution, such an analysis for Megatheriidae and other ancient Xenarthrans will become possible in future work as additional ancient nuclear genomes are made available.

### Conclusions

While extant Xenarthrans are small bodied, recently extinct Xenarthrans such as *Megatherium* were among the largest known land mammals indicating they must have had exceptional cancer suppression mechanisms. We found that intrinsic cancer risk was dramatically reduced in the stem-lineages of Xenarthra, Pilosa, and Cingulata, as well as several extinct large Xenarthran species, coincident with bursts of duplication of tumor suppressor genes. These genes were enriched in pathways that regulate apoptosis and the cell cycle which may be related to the evolution of anti-cancer cellular phenotypes in the Xenarthran lineage. We found that sloth, armadillo, and anteater cells, for example, were very sensitive to DNA damage induced by the genotoxic drug mitomycin C; repair of mitomycin C induced DNA damage involves multiple pathways such as nucleotide excision repair, homologous recombination repair and translesion bypass pathways (Lee et al., 2006), suggesting these pathways have evolved Xenarthran-specific functions. Pathways related to the cell cycle were also enriched among genes that duplicated in the stem-lineage of Xenarthra and Pilosa and cell cycle dynamics as measured by doubling times of cells in culture are longer in sloths compared to all other mammals. Finally our data suggest that armadillos, which have been the sole animal model of leprosy (Adams et al., 2012; Kirchheimer and Storrs, 1971; Storrs et al., 1975), have particularly low cancer prevalence and apparent lack of natural metastatic cancer despite the ability to experimentally induce metastasis (Marin-Padilla and Benirschke, 1963), are also a promising model organism to study the mechanisms that underlie cancer initiation and progression.

## Materials and Methods

### Ancestral Body Size Reconstruction

We first assembled a time-calibrated supertree of Eutherian mammals by combining the time-calibrated molecular phylogeny of Bininda-Emonds *et al*. (Bininda-Emonds et al., 2007), the time-calibrated total evidence Afrotherian phylogeny from Puttick and Thomas (Puttick and Thomas, 2015) and the time-calibrated Xenarthran phylogeny of Delsuc *et al*. (Delsuc et al., 2016). To construct a Eutherian supertree we replaced the Afrotherian clade in the Bininda-Emonds *et al*. phylogeny with the Afrotherian phylogeny of Puttick and Thomas and the Xenarthran phylogeny Delsuc et al (2019) using Mesquite. Next, we jointly estimated rates of body mass evolution and reconstructed ancestral states using a generalization of the Brownian motion model that relaxes assumptions of neutrality and gradualism by considering increments to evolving characters to be drawn from a heavy-tailed stable distribution (the “Stable Model”) implemented in StableTraits (Elliot and Mooers, 2014). The stable model allows for large jumps in traits and has previously been shown to out-perform other models of body mass evolution, including standard Brownian motion models, Ornstein–Uhlenbeck models, early burst maximum likelihood models, and heterogeneous multi-rate models (Elliot and Mooers, 2014).

### Estimating the Evolution of Cancer Risk

The dramatic increase in body mass and lifespan in some *Afrotherian* lineages, and the relatively constant rate of cancer across species of diverse body sizes, indicates that those lineages must have also evolved reduced cancer risk. To infer the magnitude of these reductions we estimated differences in intrinsic cancer risk across extant and ancestral *Afrotherians*. Following Peto (Peto, 2015), we estimate the intrinsic cancer risk (*K*) as the product of risk associated with body mass and lifespan. To determine (*K*) across species and at ancestral nodes (see below), we first estimated ancestral lifespans at each node. We used Phylogenetic Generalized Least-Square Regression (PGLS), using a Brownian covariance matrix as implemented in the R package *ape* (Paradis et al., 2004), to calculate estimated ancestral lifespans using our estimates for body size at each node. To estimate the intrinsic cancer risk of a species, we first inferred lifespans at ancestral nodes using PGLS and the model:

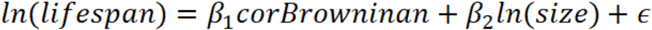

Next, we calculated *K*_1_*K*_1_ at all nodes, and then estimated the fold-change in cancer susceptibility between ancestral and descendant nodes. Next, in order to calculate *K*_1_ at all nodes, we used a simplified multistage cancer risk model for body size *D* and lifespan *t: K* ≈ *Dt*^6^ (Peto, 2015). The fold change in cancer risk between a node and its ancestor was then defined as 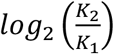.

### BUSCO assessment of genome assembly and annotation completene

We used BUSCO (Manni et al., 2021)(Galaxy Version 4.1.4) to assess genome assembly and annotation completeness of publicly available Xenarthran genomes, including Nine-banded armadillo (*Dasypus novemcinctus:* dasNov3), Screaming hairy armadillo (*Chaetophractus vellerosus*: ChaVel_v1_BIUU), Southern three-banded armadillo (*Tolypeutes matacus*: TolMat_v1_BIUU), Hoffman’s two-toed sloth (*Choloepus hoffmannii*: C_hoffmannii-2.0.1_DNAZoo_HiC), Linnaeus’s two-toed sloth (*Choloepus didactylus*: ChoDid_v1_BIUU), Brown-throated three-toed sloth (*Bradypus variegatus*: BraVar_v1_BIUU), Southern tamandua (*Tamandua tetradactyla*: TamTet_v1_BIUU), Giant anteater (*Myrmecophaga tridactyla*: MyrTri_v1_BIUU). BUSCO was run using Mode=Genome and Lineage=Eukaryota. We assessed genome quality and completeness using core Eukaryota genes because this gene set is the most conserved and thus provides a robust estimation of completeness to guide our methods to identify duplicate genes (**Figure 5**).

### Identification of gene duplications

We previously developed a reciprocal best hit BLAT (RBHB) pipeline to identify putative homologs and estimate gene copy number across species. The RBHB strategy to identify duplicate genes is conceptually straightforward: 1) Given a gene of interest *G*_*A*_ in a query genome *AA*, one searches a target genome *B* for all possible matches to *G*_*A*_ *G*_*A*_; 2) For each of these hits, one then performs the reciprocal search in the original query genome to identify the highest-scoring hit; 3) A hit in genome *B* is defined as a homolog of gene *G*_*A*_ if and only if the original gene *G*_*A*_ *G*_*A*_ is the top reciprocal search hit in genome *AA*. We selected BLAT as our algorithm of choice, as this algorithm is sensitive to highly similar (>90% identity) sequences, thus identifying the highest-confidence homologs while minimizing many-to-one mapping problems when searching for multiple genes. RBHB performs similar to other more complex methods of orthology prediction and is particularly good at identifying incomplete genes that may be fragmented in low quality/poorly assembled regions of the genome (Hernández-Salmerón and Moreno-Hagelsieb, 2020; Johnson, 2007; Kent, 2002; Vazquez and Lynch, 2021; Ward and Moreno-Hagelsieb, 2014)

In fragmented genomes, many genes will be split across multiple scaffolds, which results in BLA(S)T-like methods calling multiple hits when in reality there is only one gene. To compensate for this, we developed a novel statistic, Estimated Copy Number by Coverage (ECNC), which averages the number of times we hit each nucleotide of a query sequence in a target genome over the total number of nucleotides of the query sequence found overall in each target genome (Vazquez and Lynch, 2021). This allows us to correct for genes that have been fragmented across incomplete genomes, while accounting for missing sequences from the human query in the target genome. Mathematically, this can be written as:

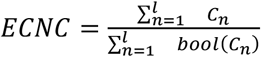

Where *n* is a given nucleotide in the query, *l* is the total length of the query, *C*_*n*_ is the number of instances that *n* is present within a reciprocal best hit, and bool (*C*_*n*_) is 1 if *C*_*n*_ >1*C*_*n*_ > 0 or 0 if *C*_*n*_ =1*C*_*n*_ = 0.

Our assessments of genome quality using BUSCO indicate that highest quality genomes were Nine-banded armadillo (*Dasypus novemcinctus:* dasNov3) and Hoffman’s two-toed sloth (*Choloepus hoffmannii*: C_hoffmannii-2.0.1_DNAZoo_HiC), while the other genomes were of much lower quality as assessed by the large number of missing BUSCOS which was 20 and 31 for Nine-banded armadillo (*Dasypus novemcinctus:* dasNov3) and Hoffman’s two-toed sloth (*Choloepus hoffmannii*: C_hoffmannii-2.0.1_DNAZoo_HiC), respectively, and ranged from 65-159 for the other genomes (**Figure 5**). These data suggest that computational inferences of copy number variation will lead to many incorrect inferences of gene duplication and loss in the lower quality genomes. Therefore, we used the RBHB/ECNC pipeline to computationally identify gene duplications in the high-quality Nine-banded armadillo (*Dasypus novemcinctus:* dasNov3) and Hoffman’s two-toed sloth (*Choloepus hoffmannii*: C_hoffmannii-2.0.1_DNAZoo_HiC) genomes, and cross-referenced this set of gene duplications with Ensembl inferred same-species paralogies (within_species_paralog) to identify a “high-quality” set of duplicated genes (Altenhoff and Dessimoz, 2009), i.e. genes inferred as duplicated with multiple methods.

Next, we used Ensembl BioMart to download the cDNA sequences for the “high-quality” duplicated genes from Nine-banded armadillo (*Dasypus novemcinctus:* dasNov3) and Hoffman’s two-toed sloth (*Choloepus hoffmannii*: choHof1), and used BLAST (discontiguous megablast) to manually determine the copy number of these genes in the lower quality Screaming hairy armadillo (*Chaetophractus vellerosus*: ChaVel_v1_BIUU), Southern three-banded armadillo (*Tolypeutes matacus*: TolMat_v1_BIUU), Hoffman’s two-toed sloth (*Choloepus hoffmannii*: C_hoffmannii-2.0.1_DNAZoo_HiC), Linnaeus’s two-toed sloth (*Choloepus didactylus*: ChoDid_v1_BIUU), Brown-throated three-toed sloth (*Bradypus variegatus*: BraVar_v1_BIUU), Southern tamandua (*Tamandua tetradactyla*: TamTet_v1_BIUU), and Giant anteater (*Myrmecophaga tridactyla*: MyrTri_v1_BIUU) genomes. Note that this approach allows us to determine if “high-quality” duplicated genes in the Nine-banded armadillo (*Dasypus novemcinctus:* dasNov3) and Hoffman’s two-toed sloth (*Choloepus hoffmannii*: choHof1) are present (ancestral) in Xenarthran lineages but cannot identify gene duplications unique (derived) in those lineages.

### RecSearch Pipeline

We previously described a Python pipeline for automating RBHB searches between a single reference genome and multiple target genomes using a list of query sequences from the reference genome (Vazquez and Lynch 2021). For the query sequences in our search, we used the hg38 UniProt proteome [*38*], which is a comprehensive set of protein sequences curated from a combination of predicted and validated protein sequences generated by the UniProt Consortium. Next, we excluded genes from downstream analyses for which assignment of homology was uncertain, including uncharacterized ORFs (991 genes), LOC (63 genes), HLA genes (402 genes), replication dependent histones (72 genes), odorant receptors (499 genes), ribosomal proteins (410 genes), zinc finger transcription factors (1983 genes), viral and repetitive-element-associated proteins (82 genes) “Uncharacterized,” “Putative,” or “Fragment” proteins (30724 genes), leaving a final set of 37,582 query protein isoforms, corresponding to 18,011 genes. We then searched for all copies of 18011 query genes in publicly available Xenarthran genomes using the RBBH/ECNC approach described above (Vazquez and Lynch, 2021).

### Reconstruction of Ancestral Copy Numbers

We encoded the copy number of each gene for each species as a discrete trait ranging as 0 (one gene copy) or 1 (duplicated) and used IQ-TREE2 (Minh et al., 2020) and ModelFinder (Kalyaanamoorthy et al., 2017) to select the best-fitting model of character evolution, which was inferred to be a General Time Reversible type model for morphological data (GTR2) with empirical character state frequencies (F0). Gene duplication events across the phylogeny were identified with the empirical Bayesian ancestral state reconstruction (ASR) method implemented in IQ-TREE2 (Minh et al., 2020), the best fitting model of character evolution (GTR2+F0), and the unrooted species tree for *Atlantogenata*. We considered ancestral state reconstructions to be reliable if they had Bayesian Posterior Probability (BPP) ≥ 0.80; less reliable reconstructions were excluded from pathway analyses. Note that in Vazquez and Lynch (2022), copy number states were coded as multi-state data corresponding to the actual copy number of each duplicate, however, many Xenarthran genomes are relatively low quality. Thus, while we identify duplicate genes in each genome the inference of actual copy number is unreliable for some species. Therefore, we used binary coding.

### Pathway Enrichment Analysis

To determine if gene duplications were enriched in particular biological pathways, we used the WEB-based Gene SeT AnaLysis Toolkit (WebGestalt) to perform Over-Representation Analysis (ORA) using the Reactome database (Jassal et al., 2020; Liao et al., 2019). Gene duplicates in each lineage were used as the foreground gene set, and the initial query set was used as the background gene set. WebGestalt uses a hypergeometric test for statistical significance of pathway over-representation, which we refined using the Benjamini-Hochberg FDR multiple-testing correction in WebGestalt.

### Cell culture, doubling time estimation, and ApoTox-Glo assay

Primary dermal fibroblasts from African elephant (*Loxodonta africana*), Asian elephant (*Elephas maximus*), South African rock hyrax (*Procavia capensis*), East African aardvark (*Orycteropus afer*), Southern Three-banded armadillo (*Tolypeutus matacus*; female, passage 6), screaming hairy armadillo (*Chaetophractus vellerosus*; female, passage 6), six-banded armadillo (*Euphractus sexcinctus*; female, passage 6), Northern tamandua (*Tamandua mexicana*; male, passage 6), Southern tamandua (*Tamandua tetradactyla tetradactyla*; female, passage 6), Linnaeus’s two-toed sloth (*Choloepus didactylus*; male, passage 6), Hoffmann’s two-toed sloth (*Choloepus hoffmanni*; male, passage 10), human, bat *(Myotis lucifugus*, passage 14), turtle (*Terrapene carolina*, passage 14), and chicken (*Gallus gallus*, passage unknown) were maintained in T-75 culture flasks in a humidified incubatopn chamber at 37°C with 5% CO_2_ in a culture medium consisting of FGM/EMEM (1:1) supplemented with insulin, FGF, 10% FBS and Gentamicin/Amphotericin B (FGM-2, singlequots, Clonetics/Lonza); cells were divided before reaching 80% confluency. For drug treatments, 10,000 cells were seeded into each well of an opaque bottomed 96-well plate, leaving a row with no cells (background control) and treated 12 hours after seeding with a serial dilution of Mitomycin C (0 uM, 5 uM, 10 uM, 15 uM, 20 uM, and 25 uM), and 12 biological replicates for each condition. After 12 hrs of incubation cell viability, cytotoxicity, and caspase-3/7 activity were measured using the ApoTox-Glo Triplex Assay (Promega) in a GloMax-Multi+ Reader (Promega). Data were standardized to no-drug (0 uM) control cells. For doubling time estimation, cells were harvested from T75 flasks at 80% confluency and seeded into six-well culture plates at 10,000 cells/well. Cell counts were performed automatically using the confluence estimation method using a Cytation-5 multi-mode plate reader (BioTek), doubling time was estimated as the number of hours cells took to grow from ∼20% to ∼40% confluency.

### Mammalian population doubling time estimation

We gathered population doubling times from previously published studies (Burkard et al., 2019, 2015; Carvan et al., 1994; Chen et al., 2009; Gomes et al., 2011; Mancia et al., 2012; Rajput et al., 2018; Seluanov et al., 2008; Wang et al., 2011, 2021; Wise et al., 2011, 2008; Yajing et al., 2018). Some studies did not report doubling times, but showed graphs of doubling times. For these datasets, doubling times were extracted from figures using WebPlotDigitizer Version 4.5 (Rohatgi, 2021).

### Cancer data collection from NHDP nine-banded armadillo

Following Aktipis et al. (2015), we defined any neoplastic growth as a cancer. The NHDP nine-banded armadillo (*Dasypus novemcinctus*) colony is comprised of both wild-caught adult and captive-born sibling animals. The wild-caught adults, including pregnant females that subsequently gave birth to genetically identical quadruplets in captivity, were collected from various locations in Louisiana and Arkansas. Adult and juvenile animals were housed singly or in pairs in modified rabbit cages with soft plastic flooring that are ganged together with a tunnel to separate the sleeping and feeding area from litter pan area. The wild-caught adults received the following treatment upon entry into the colony: Penicillin G Procain (2 mL IM) repeated at 5 days, Praziquantel (0.4 mL IM at day 7), Ivermectin (0.1 ml in the food at day 14) and Prednisone 0.25 mL IM at day 21). The captive-born siblings were treated with Fenbendazole at ∼ 6-10 weeks of age. At ∼16 months of age, the animals were treated with Penicillin (1.0 mL) and dewormed with Ivermectin (0.1 mL) and Praziquantel (0.4 mL). Prednisone (10mg/mL) was also given at this time. All animals were conditioned to captivity for ∼1year prior to experimentation.

To determine their immune response to *Mycobacterium leprae*, armadillos were injected intradermally with lepromin (heat-killed nude mice footpad derived M. leprae strain Thai 53). At 21 days, the sites were biopsied using a 4 mm punch biopsy and examined histopathologically. Armadillos with a lepromatous leprosy response were infected intravenously in the saphenous vein with 1 × 109 viable *M. leprae* derived from athymic nude mice. The NHDP armadillo colony is generally composed of ∼60% M. leprae-infected armadillos and ∼40% naïve armadillos at any given time. Experimental inoculations are done every 4 months; therefore, there are armadillos progressing at different leprosy stages throughout the colony.

Tissues were collected from the *M. leprae*-infected armadillos and used for the production of leprosy research reagents (Larsen et al., 2020). Four hours prior to sacrifice, the animal was given Gentamicin and Penicillin. The armadillo was anesthetized using a combination of Ketamine (0.6 mL) and Dexdomitor (0.4 mL) given IM, and the skin of the abdomen was shaved and cleaned with iodine, 70% ethanol, and sterile water. The animal was euthanized by exsanguination and placed on its back in a BSC where the tissues were aseptically removed. The tissues were placed in sterile jars and, after checking for contaminants, stored at 70°C.

### Cancer data collection from previous studies

Published cancer prevalence data for from giant anteater (*Myrmecophaga tridactyla*), southern tamandua (*Tamandua tetradactyla*), silky anteater (*Cyclopes* sp.), maned three-toed sloths (*Bradypus torquatus*), brown-throated sloths (*Bradypus variegatus*), pale-throated sloth (*Bradypus tridactylus*), Linné’s two-toed sloth (*Choloepus didactylus*), three-toed sloths (*Bradypus sp*.), and two-toed slots (*Choloepus sp*.) from four published surveys of morbidity and mortality in these species (Arenales et al., 2020a, 2020b; Diniz and Oliveira, 1999; Diniz et al., 1995). Neoplasia prevalence in these species of Xenarthra were compared to to other Therian mammals using data from two published studies that included pathology reports from 37 (Boddy et al., 2020) and 191 (Vincze et al., 2022) species of Therian mammals, as well as three species of cetaceans (Albuquerque et al., 2018); The total dataset includes cancer prevalence data from 221 species. Neoplasia reports from *Dasypus novemcinctus* from Boddy et al. were combined with our new cancer prevalence data from the NHDP colony and *Choloepus didactylus* data from Vincze et al were combined with *Choloepus didactylus* data from (Arenales et al., 2020b). Confidence intervals (95%) on lifetime neoplasia prevalence were estimated in PropCI package (https://github.com/shearer/PropCIs) in R using the Clopper-Pearson exact CI function: exactci(x, n, conf.level=0.95), where x is the number of successes (necropsies with neoplasia), n is the total sample size (number of necropsies), and conf.level=0.95 is the 95% lower and upper confidence intervals.

## Acknowledgements

We would like to thank R. Stevenson for collecting cancer incidence data from NHDP necropsy records and the San Diego Frozen Zoo for providing Xenarthran cell lines.

