## Supplementary figures and images for "Parallel evolution of reduced cancer risk and tumor suppressor duplications in Xenarthra"

### goslim_summary_wg_result1657297622.png

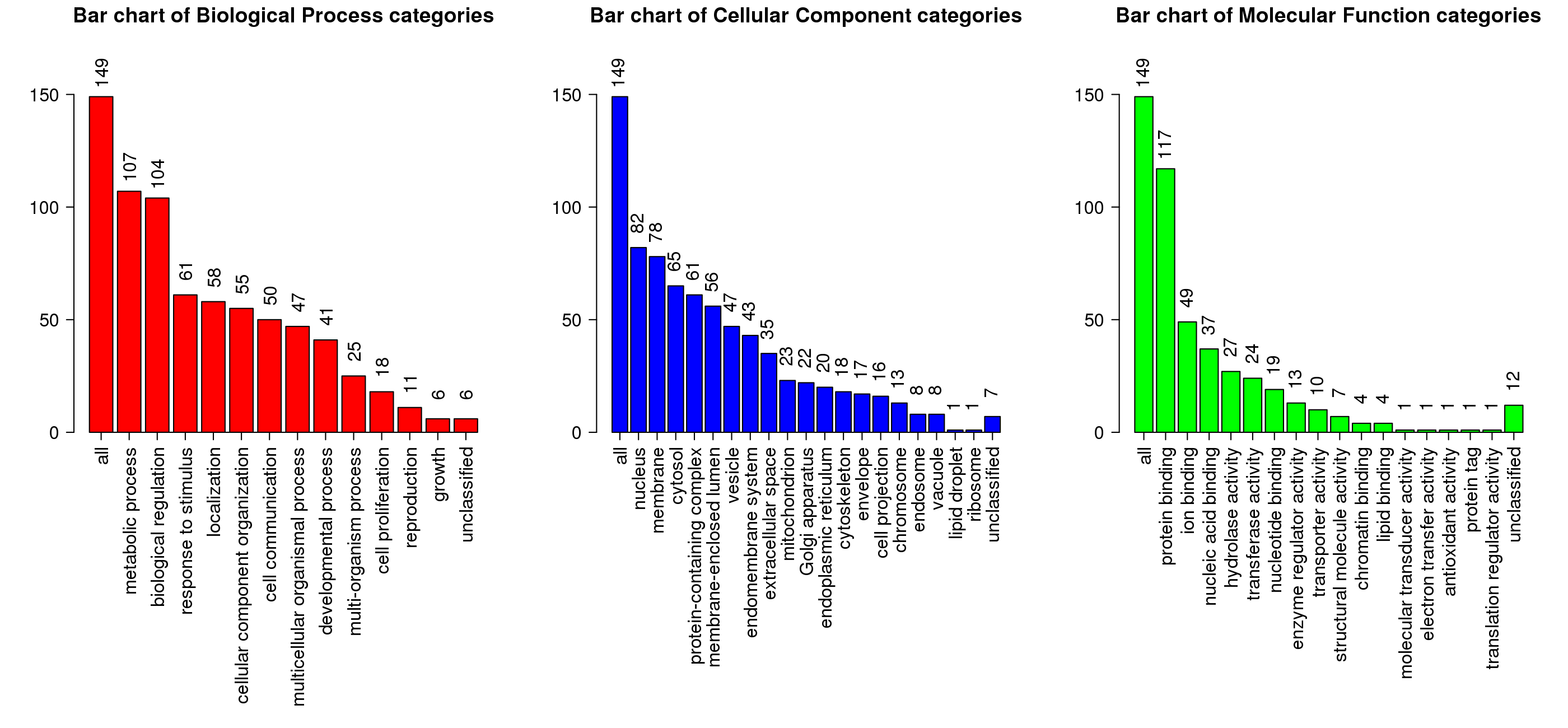

### goslim_summary_wg_result1657298103.png

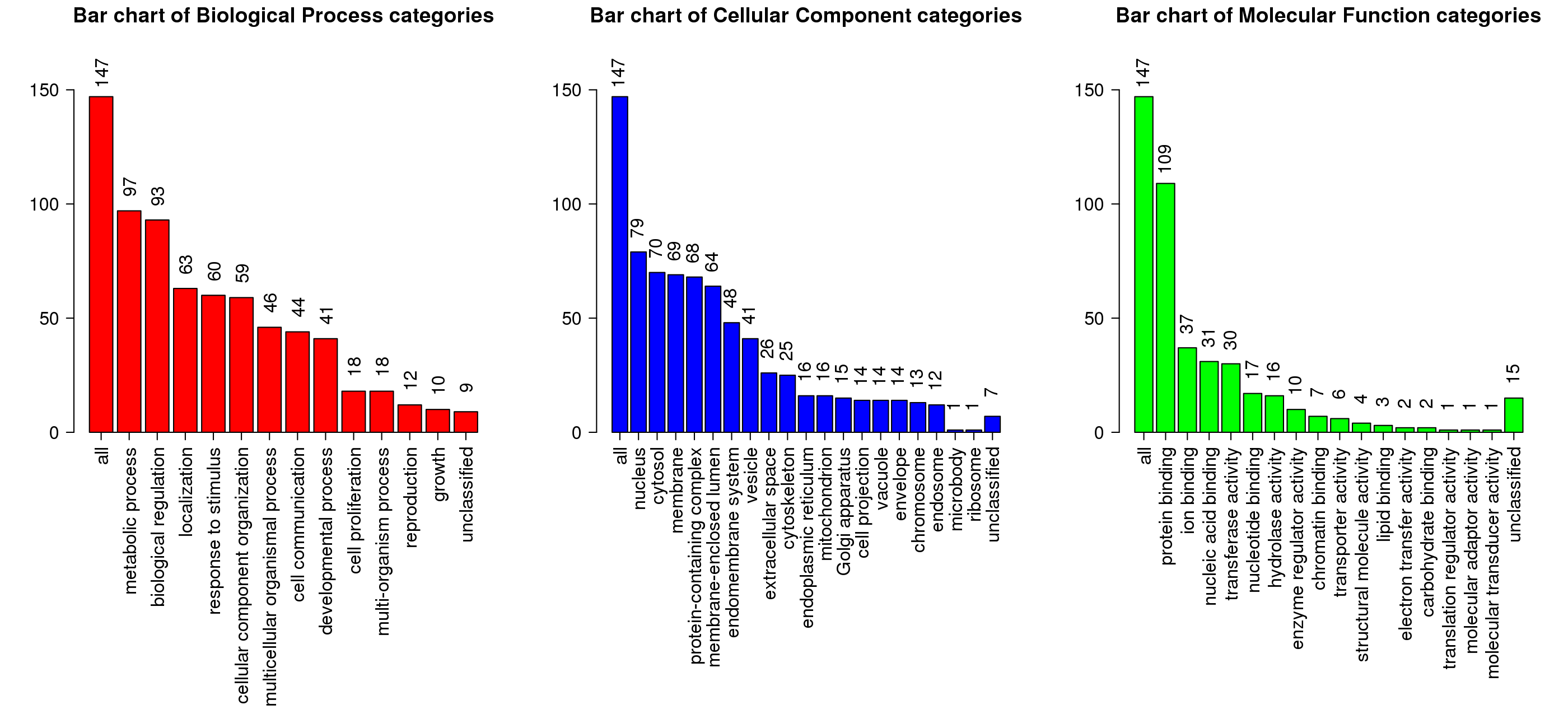

### goslim_summary_wg_result1657298133.png

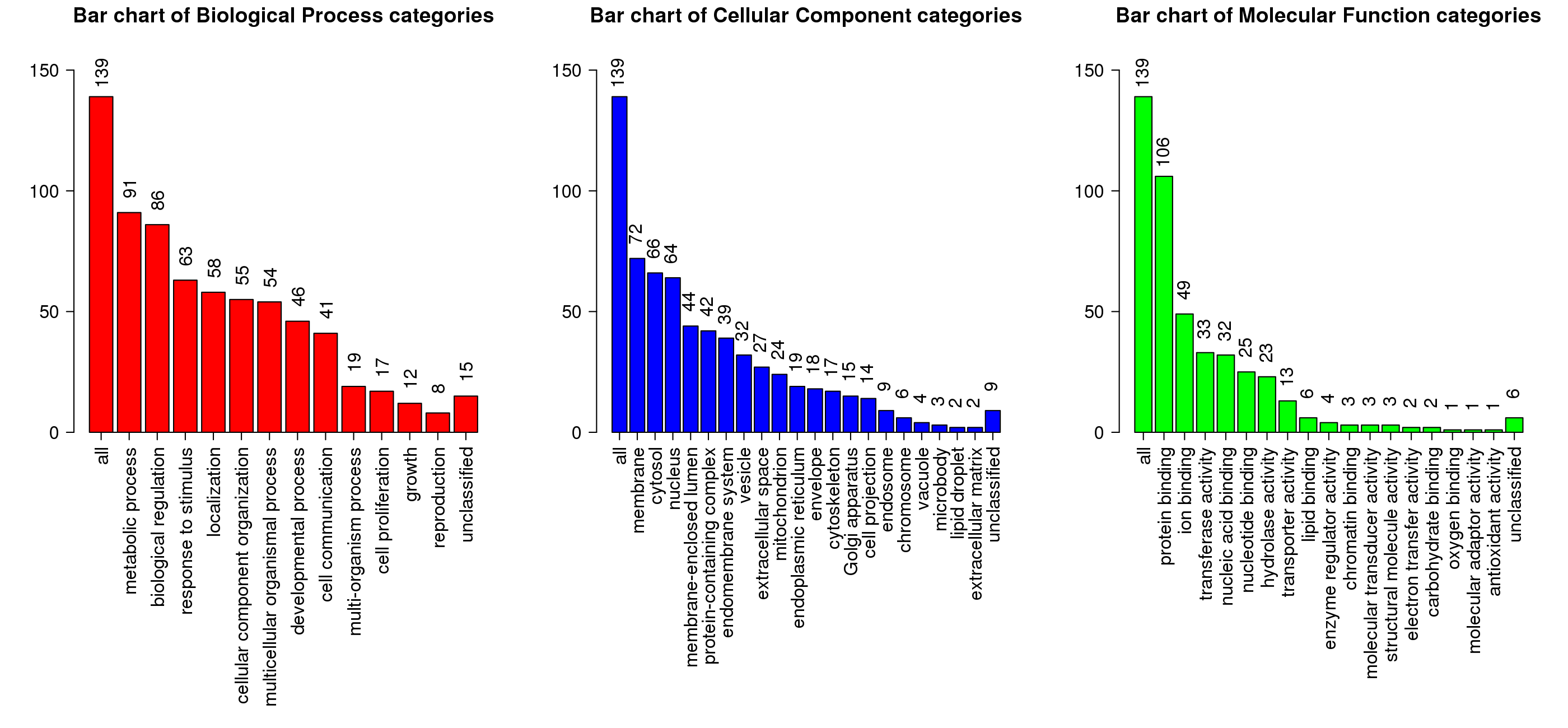
